# Patterns of urine scent mark pheromone evolution in house mice and relatives (Muridae: *Mus*)

**DOI:** 10.1101/597203

**Authors:** Michael J Sheehan, Polly Campbell, Caitlin H Miller

## Abstract

Scent marks are important mediators of territorial behavior and sexual selection in many species, especially among mammals. As such, the evolution of compounds used in scent marks has the potential to inform our understanding of signal evolution in relation to social and sexual selection. A major challenge in studies of chemical communication is that the link between semiochemical compounds and genetic changes is often unclear. The major urinary proteins (MUPs) of house mice are elaborated pheromone blends that provide information on sex, status and individual identity. Importantly, MUPs are a direct protein product of genes, providing a clear link between genotype and phenotype. Here we examine the evolution of urinary pheromone signals among house mice and relatives by examining the sequences and patterns of expression of MUPs in the liver, where urine excreted MUPs are produced. MUP patterns have evolved among mouse species both by gene duplication and variation in expression. Notably, the sex-specificity of pheromone expression that has previously been assumed to be male-specific varies considerably across species. Our data reveal that individual identity signals in MUPs evolved prior to 0.35 million years ago and have rapidly diversified through recombining a modest number of perceptually salient amino acid variants. Amino acid variants are much more common on the exterior of the protein where they could interact with vomeronasal receptors, suggesting that perception have played a major role in shaping MUP diversity. Collectively, these data provide new insights into the diverse processes and pressures shaping pheromone signals, and suggest new avenues for using house mice and their wild relatives to probe the evolution of signals and signal processing.

## INTRODUCTION

Scent marks provide complex blends of socially salient information that are important drivers of behavioral interactions for many animal species (Gosling and Roberts 2001; Hurst and Beynon 2004; Ferkin 2019). Depending on the species and source of the scent, markings may contain information about species identity, sex, age, health, physiological state, kinship, group membership, and individual identity (Wyatt 2003; Wyatt 2010). Information in scent marks may reflect the genetic makeup (Mateo 2003; Charpentier et al. 2010; Sheehan et al. 2016) as well as the behavioral and social state of an individual (Ferris et al. 1987; Martín and Lopez 2007; Freeman et al. 2019). Given the importance of scent communication for many species, there has been considerable interest in the evolutionary patterns of scent mark composition and information content across species. Pheromone diversification within and between species has been of particular interest (Fang et al. 2009; Lassance et al. 2010; Janssenswillen et al. 2014; Tupec et al. 2019) because pheromones mediate social and sexual interactions within species, and potentially act as a pre-zygotic barriers between populations (Ganem et al. 2014).

House mice are highly territorial scent markers. As the principal mammalian model organism (Phifer-Rixey and Nachman 2015), house mice are an important model for understanding the structure and function of scent marks (Desjardins et al. 1973; Hurst 1987; Thonhauser et al. 2013). Both laboratory and wild house mice use urine to mark territories. These territorial urine marks are subsequently used to assess competitors and potential mates. Work on genetically diverse wild populations of house mice has consistently demonstrated a critical role for major urinary proteins (MUPs) in scent marking (Hurst and Beynon 2013; Kaur et al. 2014). MUPs are involatile lipocalin proteins that act directly as pheromones, and also influence volatile components of urine markings (Timm et al. 2001; Nevison et al. 2003). The mouse genome encodes more than 20 tandemly arrayed *Mup* genes, a subset of which are expressed highly in the liver and excreted in the urine at high concentrations (Logan et al. 2008; Mudge et al. 2008). House mice produce two different classes of urinary MUPs with distinct expression profiles. The first class of urinary MUPs are known as central MUPs, due to their physical position within the tandem gene array. Males and females express a diversity of highly similar central MUP isoforms. While the total amount of central MUPs produced varies with social conditions (Nelson et al. 2015), the specific pattern of MUPs is genetically determined and fixed for a given individual (Sheehan et al. 2016). Importantly, the number and identity of central MUPs varies among individuals within a population, allowing MUP signatures to serve as a signal of individual identity (Sheehan et al. 2016). Individual recognition of scent marks via central MUPs is used in male-male competition, female assessment of males, and for kin recognition (Ramm et al. 2008; Hurst 2009). In addition to their role in individual recognition, variation in central MUPs allows mice to avoid mating with close relatives, cooperate with kin, and assess heterozygosity (Thom et al. 2008; Green et al. 2015). The second class of urinary MUPs, Mup3 and Mup20, have been characterized as male-specific pheromones in common lab strains (Mudge et al. 2008; Roberts et al. 2010). Due to their sex-biased expression, these MUPs appear to indicate whether a scent mark has been made by a male or female. Mup3 has been shown to stimulate male-male countermarking and aggression (Kaur et al. 2014). Mup20 (also known as darcin) stimulates learning about spatial locations of male scent marks in females and promotes countermarking in males (Roberts et al. 2012; Kaur et al. 2014).

The MUP signatures in wild mouse populations are pheromone blends that provide important social information on individual identity and sex. Previous analyses of mammalian genomes and rodent urine have shown that the patterns of MUP production seen in house mice are not widespread among rodents (Beynon et al. 2008; Logan et al. 2008; Hagemeyer et al. 2011). This raises interesting questions about the evolutionary origins of information content in mouse MUP signatures. Specifically, we address 4 questions:

### (1) What information is encoded by MUP blends?

Current evidence suggests that MUP blends encode information about individual identity, sex, and male dominance status. Information content of signals is determined by the distribution of variation among individuals within and between groups (Dale 2006; Tibbetts et al. 2017). Individual identity information is encoded by the diverse central MUPs, genes that differ among individuals in both sequence and relative expression patterns (Hurst et al. 2001; Kaur et al. 2014; Sheehan et al. 2016). Sex and male dominance status are encoded by the differential expression of *Mup3* and *Mup20.* Males are reported to express both genes while females do not (Mudge et al. 2008). Furthermore, higher status males increase expression of *Mup20* relative to other MUPs in their urine (Thoß et al. 2019). If pheromones are specific to a given species or population, they may also encode information about the species identity or population of origin (Mullen et al. 2007). Here we examine patterns of expression in males and females to assess the potential for MUPs to encode individual identity and sex across species. MUP signatures that encode individual identity could show either of two patterns, both contributing to individually recognizable signatures. First, individuals may differ in the amino acid sequences of proteins that are expressed. Second, individuals can also vary in the relative expression level of the same proteins. Similarly, MUP signatures encoding sex should show a bias in relative expression patterns between sexes of the same species.

### (2) Do specific proteins retain the same information across species?

In order for protein signatures to encode relevant social or sexual information there needs to be variation in expression among individuals or sexes. However, which proteins vary need not be the same. For example, male biased expression of a protein in one species versus female biased expression of the same protein in another, both provide information on sex. However, the information encoded by the presence of the protein differs between species. Indeed, work in *Drosophila* has shown that sex differences in expression of key pheromones differs across species, such that the information content of a particular molecule varies across the phylogeny (Seeholzer et al. 2018).

### (3) Are pheromones shared or species-specific?

In addition to encoding sex and individual information, MUPs in scent marks may encode species identity through two mechanisms (Symonds and Elgar 2008). First, the same pheromone compounds may be used in distinct ratios or blends to indicate species. Alternatively, amino acid differences between proteins may allow differential detection and response to otherwise similar protein ratios. Pheromones that are shared across species may indicate functional constraints on either MUP structure or the structure of receptors. Different pheromone compounds across species may indicate selection for divergent protein pheromone repertoires across species. Because many semiochemicals are the products of multiple genes (Lassance et al. 2010; Lassance et al. 2013), linking divergence in phenotype to function changes in genotype is a significant challenge. In the case of the mice, however, MUPs are direct gene products allowing for clear assessment of protein pheromones’ similarity from transcriptomic sequences.

### (4) What are the long-term consequences of selection for individual identity signatures?

Theory and experiments suggests that traits under selection for individuality should be under negative frequency-dependent as rare phenotypes are more recognizable and therefore facilitate correct identification (Johnstone 1997; Dale et al. 2001; Sheehan and Tibbetts 2009; Sheehan et al. 2017). While much of individuality in scent marks has been assumed to be the consequence of broadscale genetic and physiological differences among individuals (Todrank and Heth 2003; Willse et al. 2006), results from mice indicate that MUPs specifically encode individual identity (Hurst et al. 2001; Kaur et al. 2014; Roberts et al. 2018). Indeed, molecular analyses of *Mup* genes within a population of *M. m. domesticus* identified clear signatures of negative frequency-dependent selection (Sheehan et al. 2016). The longer-term consequences of selection for individual identity signatures on patterns of phenotypic diversification have yet to be explored. Two possible outcomes from negative frequency-dependent selection have been observed. On the one hand, selection may maintain particular genetic and phenotypic variants over long periods of time. For example, negative frequency-dependent selection maintains multiple male mating strategies for long periods of time in lineages such as *Uta* lizards (Sinervo et al. 2001; Corl et al. 2010). On the other hand, selection for individuality may not depend on a few common variants maintained over long periods of time, but rather on the generation of novel variants. High diversity and turnover of alleles used in some fungal mating systems are indicative of such a process (James 2012). We can begin to tease apart these possibilities by examining the extent to which different species, subspecies, or populations share particular protein variants.

Here we combine newly generated and publicly available transcriptomic data to examine the sequences and patterns of *Mup* genes expressed in the liver of house mice and close relatives. Previous work has already established that mice and rats have independently expanded their *Mup* gene families (Logan et al. 2008; Gomez-Baena et al. 2018). Therefore, we focus our efforts on species in the genus *Mus* for which liver transcriptomic sequences are available.

## METHODS

### Generation of liver transcriptomes

We harvested liver tissues from adult males and females for lab mice from multiple strains and species representing approximately 7 million years of evolution within the genus *Mus*. RNA was extracted from tissue using a Qiagen RNeasy Kit. RNA sequencing libraries were generated using the NEBNext Ultra RNA Library Prep Kit for Illumina (NEB #E7530). NEBNext Poly(A) mRNA Magnetic Isolation Module (NEB #E7490) was used for RNA Isolation. Sequences were indexed using the NEBNext Multiplex Oligos for Illumina (Dual Index Primers Set 1, NEB #E7600). We sequenced paired end libraries at the Institute for Biotechnology at Cornell University. Some libraries were sequenced on a MiSeq (PE 300) while others were sequenced on a NextSeq (PE 150). Information for specimens used in for sequencing is given in Table S1 (Supplemental File 1). All procedures were approved by the Institutional Animal Care and Use Committee of Cornell University. Newly sequenced data is available under PRJNA530260.

### Criteria for choosing publicly available liver transcriptomes for analysis

We searched the NCBI Short Read Archive (https://www.ncbi.nlm.nih.gov/sra) for liver transcriptomes of muroid rodents with the goal of identifying transcriptional data for species in *Mus* and related genera. The vast majority of sequences are derived from experiments examining differential expression in liver transcriptomes across treatments in laboratory mouse strains, which we did not consider. Rather, we focused our attention on data from unmanipulated liver samples that provided information on the expression of urinary proteins across a diverse group of mice. All told, we examined 100 samples from 10 species or subspecies (Fig 1, Table S1).

**Fig 1.**
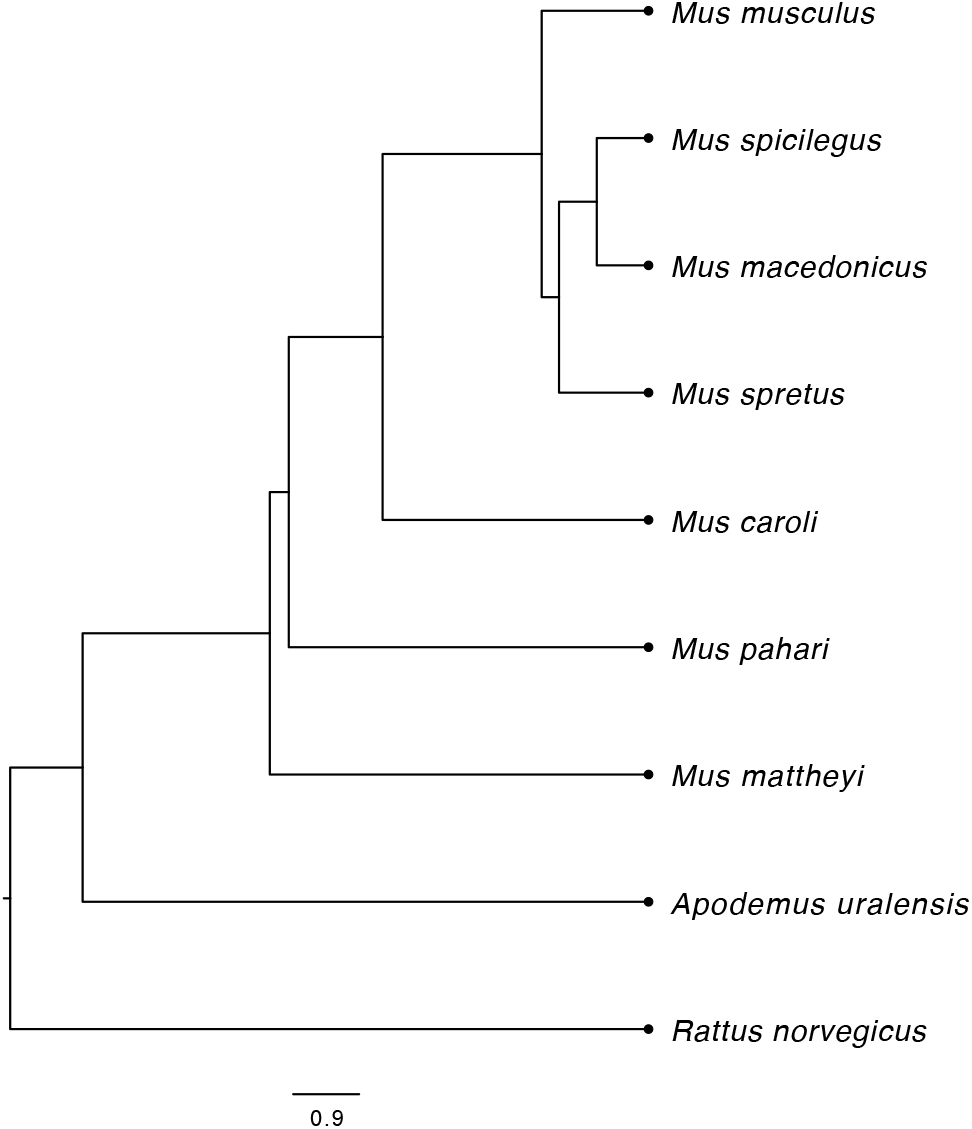
Species tree for muroid rodents examined in this study. Modified from (Steppan and Schenk 2017). The scale bar measures substitutions per site.

### *De novo* liver transcriptomes

For species that are more distantly related to the house mouse and thus more likely to have poor sequence alignment with the reference genome (*M. caroli*, *M. pahari*, *M. mattheyi and Apodemus uralensis*) we generated *de novo* liver transcriptomes to identify potential *Mup* sequences. RNAseq reads generated for this study (*M. caroli* and *M. pahari*) or downloaded from the short read archive (*M. mattheyi and A. uralensis*) were assembled using default settings in rnaSPAdes (Bushmanova et al. 2018). We then blasted known *Mup* transcript sequences from the house mouse against assembled transcriptomes to identify *Mup* transcripts. All identified transcripts appeared to be full length *Mup* genes upon manual alignment and inspection.

### Alignment of RNASeq reads

In *M. m. domesticus*, alignment of short reads from MUP genes or transcripts is problematic due to the high similarity of sequences within the central MUP cluster. Previous work examining MUP diversity (Sheehan et al. 2016) aligned sequences to a modified transcriptome consisting of sequences of *Mup3, Mup11* and *Mup20*. For distantly related species *M. caroli, M. pahari* and *M. mattheyi* we also added sequences derived from *de novo* transcriptomes. We did not detect any *Mup* genes in the transcriptome of *A. uralensis* so no sequences were added for this species. Reads were aligned with bowtie2 using the ‘–very-fast’ setting.

### Comparing patterns of gene expression

Urinary pheromone output varies among individuals both in the overall production of MUPs and the relative production of each protein within an individual (Nelson et al. 2015; Sheehan et al. 2016). In comparing gene expression our goal was to understand the relative contributions to pheromone blends of different MUP types across individuals and species. Visual inspection of read alignments to MUP transcripts showed that coverage varied along the transcripts in relation to the sequencing strategy employed. The shorter reads used in some of the Short Read Archive experiments produced more even coverage along the transcript than PE300 sequencing experiments, though longer sequences allowed for easier identification of full-length transcripts. To account for differences in sequencing strategy and depth across all the samples we scored the level of expression among genes as the deepest read count for each gene. From these measures we then calculated the relative contributions of each gene type (central *Mup* sequences aligned to *Mup11*, sequences aligned to *Mup3,* and sequences aligned *Mup20*) to composite pheromone blend for each individual. This approach has the benefit of capturing within-individual allocation of MUPs, rather than variation in MUP expression relative to other proteins in the liver as a whole, as would be the case in a typical differential expression analysis. Previous work utilizing the same approach for documenting relative *Mup* transcript abundance showed that this method predicts the final patterns of relative protein excretion in a wild population of *M. m. domesticus* at a nearly 1:1 ratio (Sheehan et al. 2016). Moreover, the tight correlation between protein excretion and liver transcripts suggests that these measures likely provide an accurate view of the relative abundance of different MUPs excreted in the urine.

### Searching for *Mup* genes in genome assemblies

We searched for *Mup* genes in published genomes for the *M. m. musculus* strain PWK/PhJ, the *M. m. castaneus* strain CAST/EiJ, and *M. spretus, M. spicilegus, M. caroli* and *M. pahari* (Keane et al. 2011; Couger et al. 2018; Thybert et al. 2018). All genomes except *M. spicilegus* were accessed via the Ensemble genome browser 95. The genome for *M. spicilegus* was downloaded from GenBank. Transcript sequences for the *Mup11, Mup3* and *Mup20* were downloaded from the Mouse Genome Informatics website (http://www.informatics.jax.org/). For all species, we blasted the transcripts of the three genes against the genome. We used the conserved exon/intron structure among *Mup* genes within the mouse genome to identify full length gene sequences in each assembly. Within the BLAST results we searched for consecutive strings of sequence covering the whole transcript over approximately 3-5kb. The entire sequences encompassing the transcript ± 1kb were then downloaded and aligned with other known *Mup* genes using the online version of MAFFT v.7 (https://mafft.cbrc.jp/alignment/software/) and then further examined by hand using Mesquite (Maddison 2008). Only sequences predicted to produce a full-length protein were retained. To classify sequences, we generated a gene tree of all predicted mRNA sequences using IQ-Tree (Nguyen et al. 2014). We generated a ML tree with 1000 bootstraps. The best evolutionary model was selected using the built-in model selection function. In this case the best mode was: TMP3+F+G4. Gene sequences identified are provided in Supplemental File 3.

### Bespoke detection of variants and assembly of full length mature MUP sequences

The high sequence similarity among central MUPs makes the automated assembly of high-confidence *Mup* gene sequences challenging. With paired end RNAseq data, however, it is possible to reconstruct the sequences of transcript sequences associated with mature MUPs (162 amino acids). Using the Integrated Genome Viewer v 2.4 (Robinson et al. 2011), we determined full-length sequences of mature MUP using the following strategy. First, we adjusted the allele cut-off for showing variants to 0.05 for central MUPs to account for the fact that in some individuals 10-20 central MUPs may be excreted. For pooled samples, this process likely misses some MUP sequences but allows for high confidence in those that are generated. For the peripheral MUPs, we kept the cut-off value at 0.3. Reads were viewed as pairs and sorted by insert size (largest inserts first). For each variant within the sequence, we sorted by base so that all variants containing a particular nucleotide were sorted together. Combinations of variants were recorded. This process was repeated for each variable site. Duplicate sequences generated by this process were noted and a final output of unique sequences for each sample was generated. Gene sequences used for analysis are provided in Supplemental File 2.

### Molecular diversity of mature central MUPs

mRNA sequences for each MUP were translated to proteins using standard RNA-DNA conversions in Mesquite. We then aligned all central MUP genes from this study as well as those previously identified from wild *M. m. domesticus* from Edmonton, Canada (Sheehan et al. 2016) to examine the distribution of amino acid variants across the proteins and their relative abundance among the proteins sampled. The 3D structure of MUP11 is a barrel of beta-sheets, characterized by an inner hydrophobic pocket that binds and transports volatile compounds, and a hydrophilic exterior that is assumed to interact with MUP-detecting vomeronasal receptors Phelan et al (2014). The model includes the relative position of each amino acid on the interior or exterior of the protein, noted as the percentage exposure to the exterior (broken into quintiles), or as part of the central barrel. Thus, we counted the number of proteins that possess a non-reference Mup11 amino acid at each position relative to how exposed it is to the exterior.

### Gene tree of *Mup* sequences

A gene tree of *Mup* transcripts corresponding to mature proteins excreted in the urine was generated from sequences aligned using the online version of MAFFT v.7 in IQTree using the same procedure as above. A single *Mup* gene sequence from the independent *Mup* gene family expansion in *Rattus* (ENSRNOT00000046760) was used as an outgroup.

## RESULTS

### Sequences of *Mup* genes expressed in the liver

Though both house mice and rats express a diversity of *Mup* genes in their livers, other mouse species and strains examined here showed wide variation in the number and types of *Mup* genes expressed (Fig 2). While we detected *Mup* genes in all *Mus* species sampled we were unable to detect any *Mup* orthologs in the liver transcriptome of *A. uralensis*, indicating that liver expression of *Mup* genes is not a universal trait among muroid rodents. Details of all of the *Mup* gene sequences and predicted proteins surveyed in this study are provided as Table S1 and S2 in Supplemental File 1.

**Fig 2.**
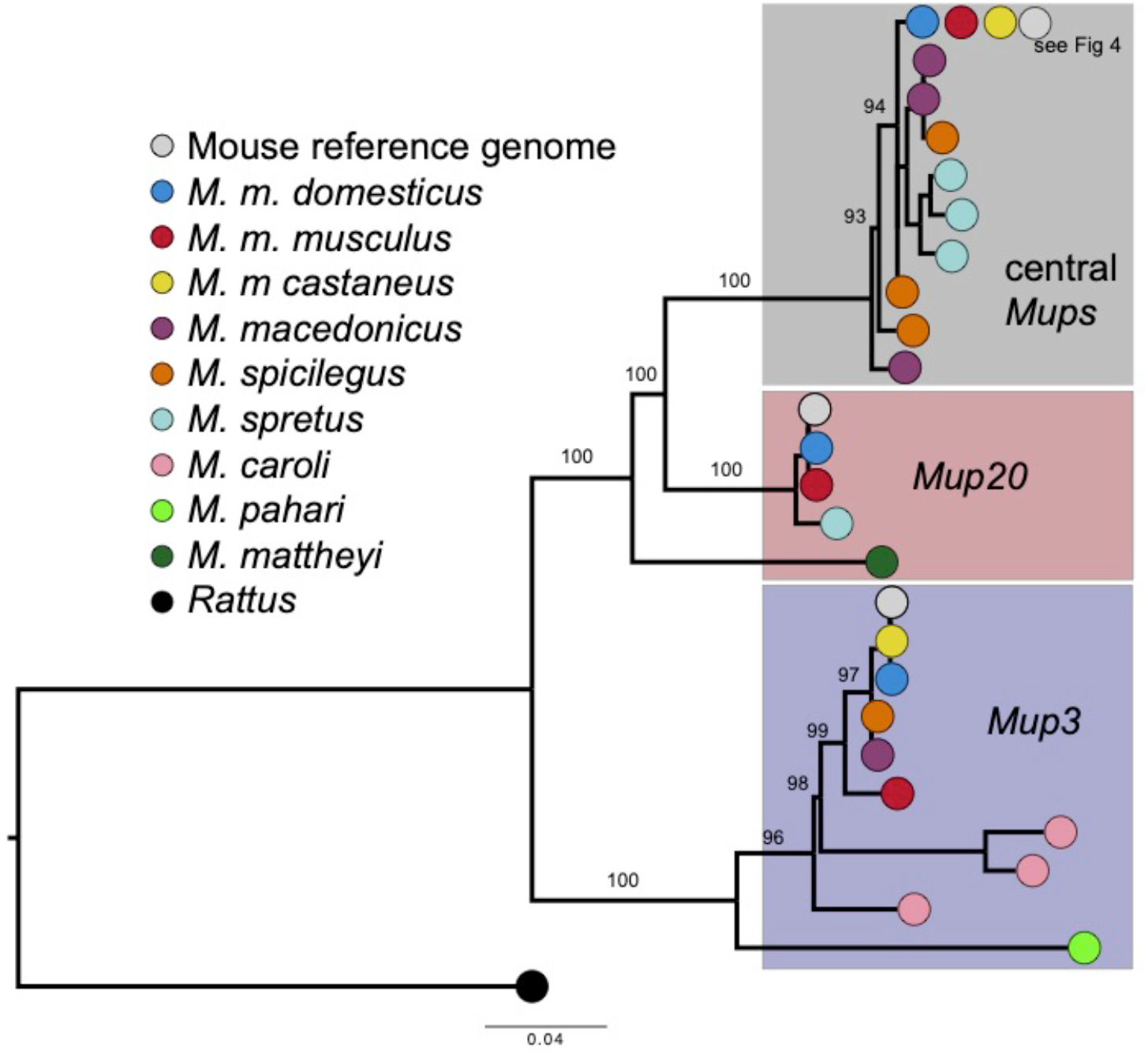
Simplified gene tree of *Mup* genes expressed in the liver. Expressed sequences for each species or subspecies are shown with a different color (see legend). Reference sequences for *Mup11, Mup3, Mup20* and an outgroup sequence for a rat MUP has been used to root the tree. For the three subspecies of *M. musculus* only data for one inbred lab strain each are shown here. The diversity of central MUPs in *M. musculus* lab and wild samples are represented here with *Mup11* sequence. See Fig 4 for additional details on central *Mup* gene diversity. The scale bar shows substitutions per site. Bootstrap values are show for major nodes. Full details in Supplemental File 4.

With the exception of the *Mup* gene expressed by *M. mattheyi,* the detected *Mup* genes can be readily classified as being central MUPs or related to Mup3 or Mup20 (Fig 2). The gene expressed by *M. mattheyi* appears to be basal to the subsequent diversification of central MUPs and *Mup20* in other species. Subspecies of *M. musculus* tend to express a single conserved version of *Mup3* and *Mup20* but other species have at least one amino acid variant differentiating sequences between species. Notably, the close relatives of house mice either express *Mup3* (*M. spicilegus* and *M. macedonicus*) or *Mup20* (*M. spretus*) but not both.

### Patterns of *Mup* gene expression in the liver vary widely across mouse species

Across species and sexes, we identified three key ways that *Mup* gene expression varies (Fig 3): (1) the level of overall investment in *Mup* gene expression, (2) which types of *Mup* genes are expressed and (3) the relative ratios of different types of *Mup* genes. The three patterns variously contribute to differences in *Mup* gene expression between sexes and among species. Combined with sequence differences in *predicted* MUPs (Fig 2), the patterns of expression suggest that MUP profiles provide information about species and sex.

**Figure 3.**
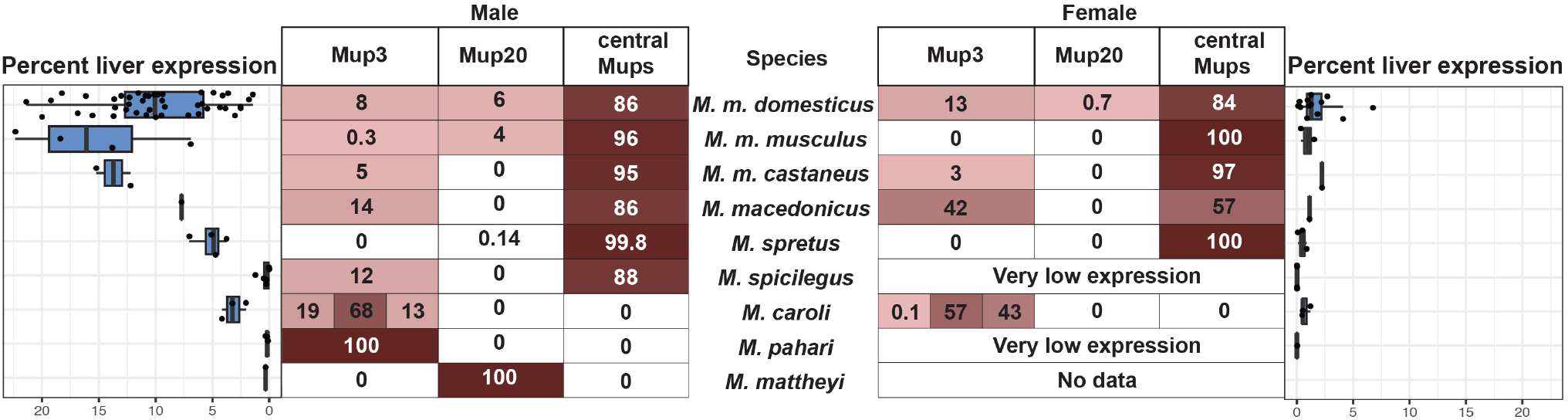
Distribution of *Mup* gene expression among species and between sexes. The average percentages of total *Mup* gene expression that is composed of *Mup3, Mup20* and central MUPs are shown for males and females of each species (where data are available). High proportions are shown as darker red. The box plots on the side of the chart show the level of total *Mup* expression, calculated as the percent of all reads that aligned to *Mup* genes, for each sample examined.

Investment in overall levels of *Mup* expression varies among species and sexes (ANOVA, species: *F_7,89_*= 10.05, *P* = 5.25e-9; sex: *F_1,89_*= 47.13, *P* = 1.17e-9, species*sex: *F_7,89_*= 2.8, *P* = 0.012). Across species, males show higher levels of *Mup* gene expression (Fig 3), consistent with patterns previously reported in subspecies of *M. musculus* (Stopkova et al. 2007). Overall levels of expression vary across species, though expression levels in males and females are correlated (linear regression, *t*_6_ = 3.16, *r^2^* = 0.63, *P* = 0.019). There is considerably higher investment in *Mup* expression in *M. musculus* subspecies males compared to other species, especially *M. spicilegus, M. pahari* and *M. mattheyi*, which have low levels of *Mup* gene expression (Fig 3). It is worth noting that even in these species with ‘low’ expression levels *Mup* genes are in fact highly expressed compared to other genes in the liver transcriptome. Levels of expression in female *M. spicilegus* and *M. pahari* are exceedingly low, however, suggesting that very little protein is likely to be found in their urine. A lack of *Mup* gene expression was also seen in male *M. spicilegus* from the wild-derived strain SPI/TUA, though males of other wild-derived *M. spicilegus* strains as well as wild-caught males expressed modest levels of *Mup* transcripts.

Patterns of expression across species and sexes revealed that *Mup3* is not a male specific gene. Female *M. m. domesticus, M. m. castaneus* and *M. macedonicus* all express Mup3 at modest levels (Fig 3). Male *M. m. musculus* express a *Mup3*-like gene at very low levels (predicted to be <<1% of total MUP content; Fig 3). This same pattern of expression is present in lab strains PWK/PhJ and CZECHII/EiJ as well as pooled samples from wild-caught male *M. m. musculus*, indicating that the low expression of an unusual *Mup3*-like gene is not an artifact of inbreeding in wild-derived strains. Our analysis also reveals that *M. caroli* has a lineage-specific expansion of *Mup3* related genes (Fig 2 and 3). Two of these genes are expressed by both males and females while a third is strongly male-biased.

With the notable exception of *M. m. domesticus*, expression of *Mup20* appears to be limited to species or subspecies that do not express *Mup3* (or only express it at very low levels). Both *M. mattheyi* and *M. spretus* express genes related to *Mup20* but not *Mup3* (Fig2 and 3). The expression of *Mup20* is very low, however, in *M. spretus*. Similarly, *M. pahari, M. caroli, M. macedonicus, M. spicilegus* and *M. m. castaneus* all express orthologs of *Mup3* but not of *Mup20*. Male *M. m. musculus* express *Mup20* at high levels (Fig 3) but, as mentioned above, express a *Mup3*-like gene at extremely low levels. In both *M. m. musculus* and *M. m. domesticus* expression of *Mup20* is strongly male-biased, consistent with previous reports (Roberts et al. 2010).

Expression of central *Mup* genes is absent *M. mattheyi, M. pahari,* and *M. caroli* but is the dominant form of *Mup* expressed in the livers of *M. spretus, M. spicilegus, M. macedonicus* and *M. musculus* subspecies (Fig 3). This pattern holds for both males and females. In fact, both *M. spretus* and *M. m. musculus* females appear to only express central *Mup* genes.

### Genomic evidence for *Mup* genes not expressed in the liver

The variation in expression of *Mup* gene types (3, 20 and central) across species could arise either because of the differential gain and loss of genes among species, through differential gene regulation, or some combination of the two. To test for the role of gene gains and losses, we examined genomic data for *M. pahari, M. caroli, M. spretus, M. spicilegus*, *M. m. musculus* (PWK/PhJ) and *M. m. castaneus* (CAST/EiJ). In all cases, the *Mup* gene cluster is poorly assembled, as is also the case for the mouse reference genome (Mudge et al. 2008). Therefore, we limited our query to whether orthologs of *Mup3, Mup20* and a central MUP could be recovered from genomic data for cases in which a transcript was not detected in liver. A maximum likelihood gene tree was estimated to determine the orthology relationships for genes identified in the examined genomes (Fig S1, Supplemental File 3).

We detected genomic evidence for genes that are not transcribed in the liver in five of the genomes examined (Table 1). We did not detect any full-length *Mup* genes in the *M. spicilegus* genome assembly, though this is likely an issue with assembly rather than a true lack of genes since we detect expression of full-length *Mup* transcripts (Fig 2 and 3). Both the *M. m. castaneus* and *M. caroli* genome assemblies contain single copy orthologs for the *Mup* genes missing from liver transcriptomes. Multiple genes in *M. pahari* were identified that are basal to the divergence of *Mup20* and central MUPs clade and may be orthologs to the liver-expressed *Mup* gene identified in *M. mattheyi*. However, weak resolution in this part of the gene tree precludes strong conclusions regarding orthology (Fig S1, Supplemental File 3). The *M. spretus* genome assembly contains evidence for multiple peripheral *Mup* genes though there is no detectable ortholog for *Mup3*. While consistent with *Mup3* loss, this finding could be an artefact of fragmentary assembly of the tandemly-arrayed *Mup* gene family.

**Table 1:** Genomic evidence for non-expressed *Mup* genes

| Species | *Mup3* | *Mup20* | central<br>MUPs |
| --- | --- | --- | --- |
| *M. m. domesticus* | T<sup>1</sup> | T | T |
| *M. m. musculus* | T | T | T |
| *M. m. castaneus* | T | Genomic | T |
| *M. spretus* | No<br>evidence | T | T |
| *M. caroli* | T | Genomic | Genomic |
| *M. pahari* | T | Genomic<sup>2</sup> |  |

(1) T = transcriptomic
(2) These genes are basal to the *Mup20* / central MUPs divergence.

### Multiple expansions of central *Mup* genes

The diversity of central *Mup* genes detected in the liver transcriptomes of *M. spretus, M. spicilegus, M. macedonicus* and *M. musculus* subspecies is shown in detail in Figure 4. Relative to the other mouse species examined here, all of these species show an expanded repertoire of central *Mup* genes, indicating that there was a gene expansion in the common ancestor of this clade. We detected three different predicted proteins expressed in *M. macedonicus* and an additional three in *M. spicilegus*, though neither forms a species-specific monophyletic clade. Previous work examined the MUPs in urine of *M. macedonicus* and found a single dominant protein with a molecular weight of 18742 DA (Robertson et al. 2007). The predicted dominant protein from the liver transcriptome is expected to have a molecular weight of 18739 DA, which is similar but outside the range of measurement error, suggesting differences between the wild samples and the wild-derived inbred lab strain. We did, however, identify a protein with mass 18742 DA in *M. spicilegus*. In two wild-derived strains, ZRU and ZBN, this single protein is the only central MUP predicted for males and it is the dominant MUP predicted for a wild-caught sample.

**Figure 4.**
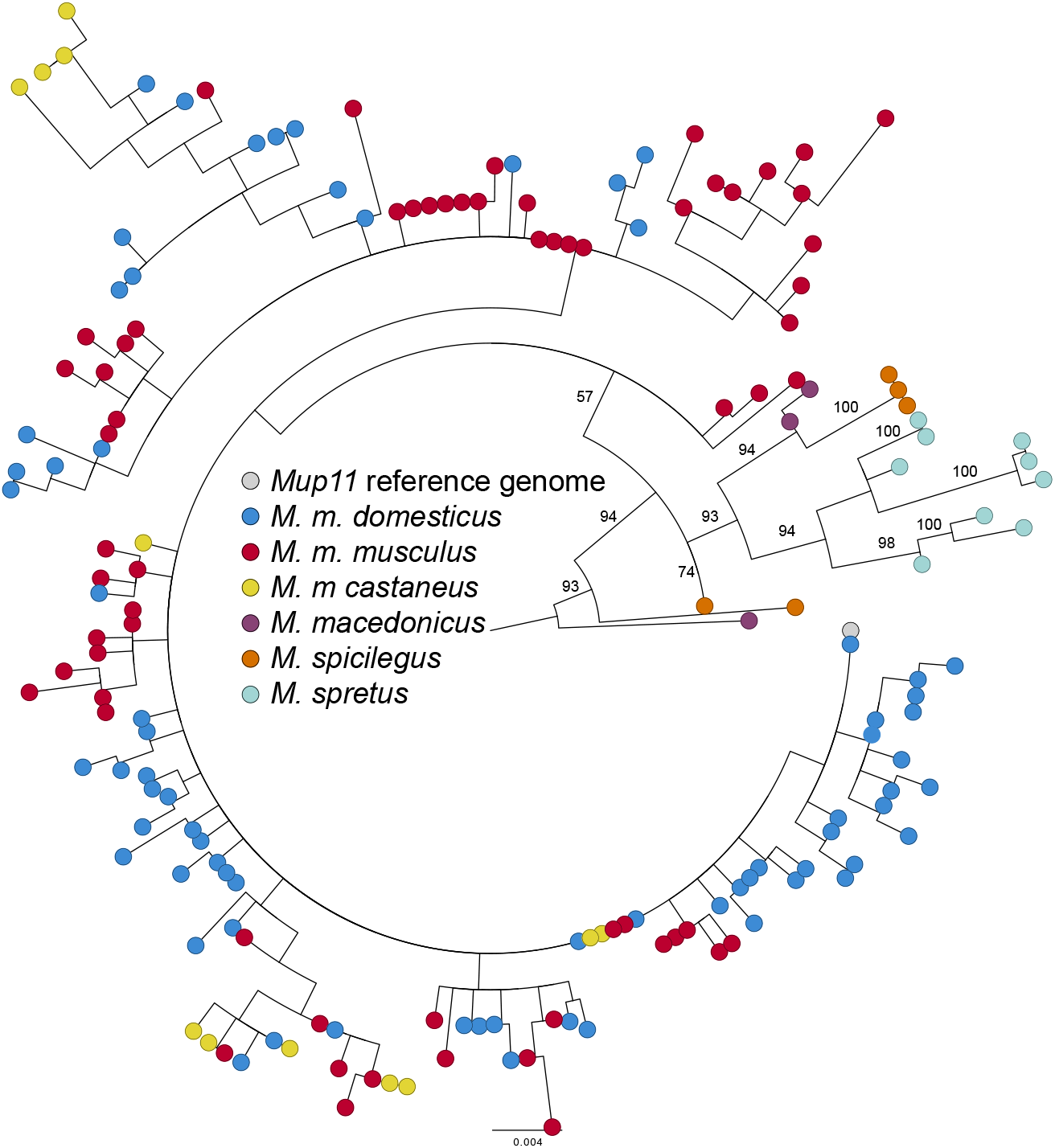
Gene tree of central *Mup* genes expressed in the livers of mice. Expressed sequences for each species or subspecies are shown with a different color (see legend). The gene tree indicates that the large expansion of central *Mup* genes previously reported for *M. m. domesticus* is shared by other *M. musculus* subspecies, suggesting that the expansion predates subspecies divergence approx. 0.35 million years ago. The scale bar shows substitutions per site. Bootstrap values are shown for major nodes. Full details for bootstrap values are provided in Supplemental File 4.

A small expansion of sequences found in wild-derived and wild-caught *M. spretus* form a monophyletic clade nested within the *M. spicilegus* and *M. macedonicus* sequences. Previous work examining the variation in MUPs in wild *M. spretus* urine identified three proteins (Beynon et al. 2008). We have identified 9 different genes sequences that are predicted to produce 6 distinct proteins. In the wild-derived strain SPRET/EiJ we found only 3 proteins. The most abundant protein has a predicted weight of 18758 DA, similar to the most abundant protein reported from wild-caught samples. The other predicted proteins from SPRET/EiJ have similar weights to previously reported proteins measured from wild *M. spretus* (18668 and 18685 versus previously reported 18666 and 18687 from Beynon et al. 2008).

The large expansion of central *Mup* sequences found among the *M. musculus* subspecies form a monophyletic group nested within the larger central *Mup* clade, although the bootstrap value of 57 indicates poor support at the node leading to *M. musculus* central *Mup* genes. The nodes within the *M. musculus* clade generally have very poor support (Supplemental file:). Weak phylogenetic signal may be caused by a combination of purifying selection on the overall gene sequences, and gene conversion (Sheehan et al. 2016).

Due to the poor resolution of the genes in the *M. musculus* clade it is difficult to draw many firm conclusions, though two patterns appear from the data. First, there are many genes found in each subspecies. This is even true for *M. m. castaneus*, represented by just two wild-derived inbred strains (CAST/EiJ and TWN). Second, parts of the expansion are shared in common across the three subspecies. One gene is shared among all three and at least two other clades feature sequences from all three subspecies. Shared expansions across all three subspecies are consistent either with a scenario in which central *Mup* genes underwent an initial expansion prior to subspecies diversification, or there has been introgression of central *Mup* genes among subspecies.

### Molecular diversity of central MUP proteins

The 165 central *Mup* transcripts detected across *M. musculus* subspecies samples (Fig 4) encode 77 distinct proteins. The 77 protein variants arise from combinations of 39 amino acid variants spread across 32 sites (Fig 5a). We examined the amino acid sequences among *M. musculus* subspecies samples and compared these to amino acid variants found in *M. spretus, M. macedonicus* and *M. spicilegus*. Only 5 of 39 amino acid variants are shared with other species. Thus, most amino acid variants appear to be specific to house mice, the majority of which were detected in only a single sample (50/77 proteins, Supplemental Table 2). Please note that this finding does **not** necessarily indicate that proteins tend to be limited to single individuals, as many of the samples examined are pooled samples. Nevertheless, it does indicate that pheromone blends composed of both rare and common proteins are likely the norm in many house mouse populations. Of the 17 proteins detected in more than one sample, 7 are shared between two subspecies and 2 are shared by all three subspecies. No proteins were shared across all samples.

**Figure 5.**
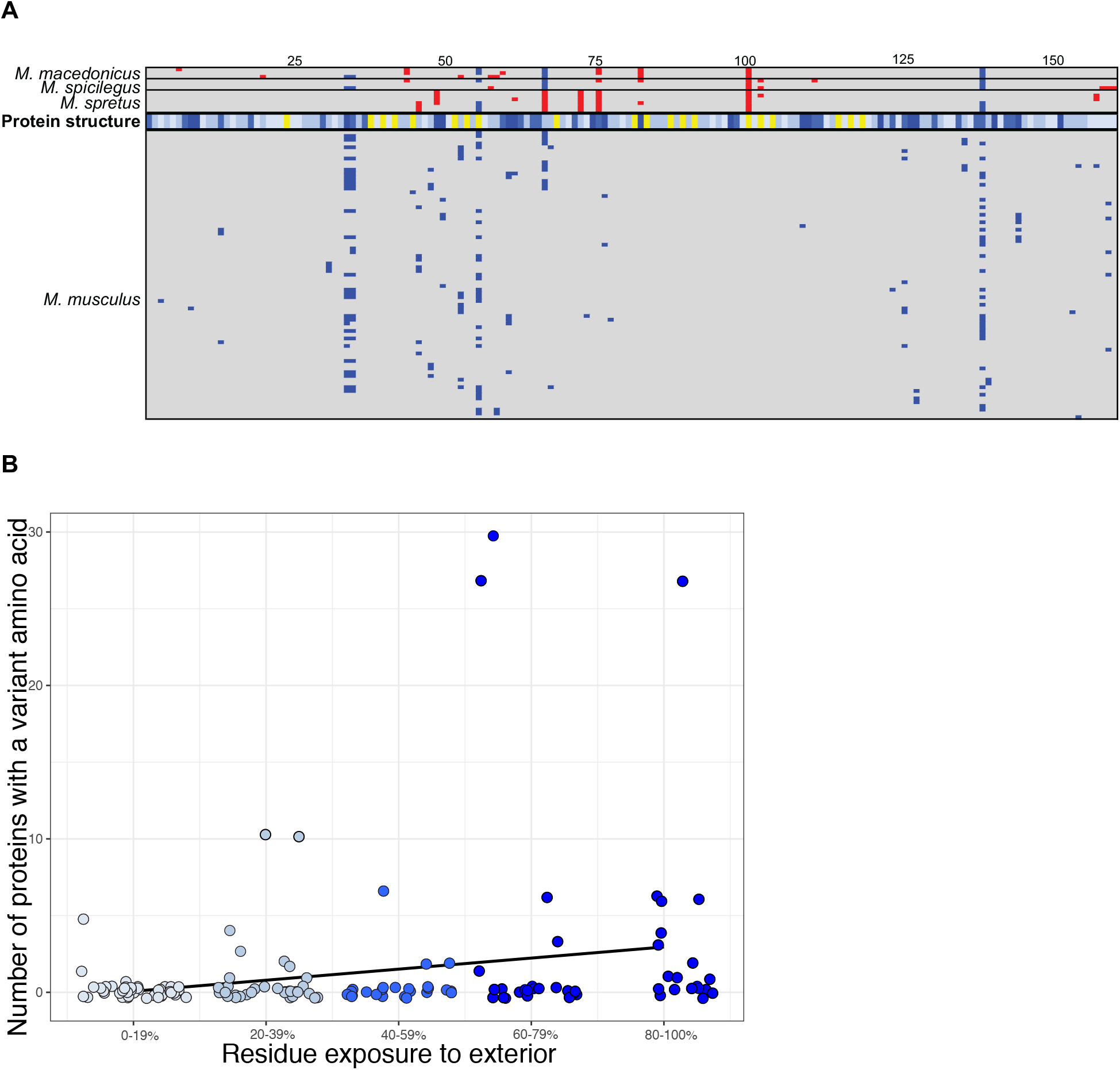
Molecular diversity of central *MUP* proteins. (A) The distribution of amino acid variants among central MUPs is depicted for mature protein sequences (162 amino acids length). Amino acid variants that differ from the sequence of a MUP11 are show in blue if found in a *M. musculus* subspecies, red if only found in a different species. Sequences for *M. spretus, M. macedonicus* and *M. spicilegus* are shown at the top while sequences for *M. musculus* subspecies are shown on the bottom. The bar labeled ‘protein structure’ indicates the position how the amino acid resides within the 3D protein structure based on Phelan et al 2014. Yellow indicates an amino acid is part of the interior beta-sheet barrel. Exposure levels on the exterior of the protein are shown as quintiles (0-20% lightest blue to 80-100% darkest blue). Greater detail, including the protein sequences, is provided in Supplemental File1. (B) Variants are more common among amino acid sites with greater exposure to the exterior of the protein, which can interact with vomeronasal receptors. The figure shows the number of protein variants containing sites with each category of exposure (excluding sites in the internal barrel). Coloring of points corresponds to the ‘protein structure’ bar in part A, where darkest blue is the highest level of exterior exposure.

We next investigated how protein structure may influence patterns of diversity. Most amino acid variants appear in one or a few proteins, though 4 variants are found in more than 25 proteins. The variable sites are predominantly found on the hydrophilic exterior surface of the protein (30/32) compared to the interior hydrophobic pocket (2/30). The extent of residue exposure predicts the level of variation observed across sites (Fig 5b, Spearman rank order correlation, *t_140_*=3.41, *r_s_*=0.27, *P*=0.0008). Thus, protein variants are more abundant at sites that likely influence binding between MUPs and vomeronasal receptors.

## DISCUSSION

Our investigation of the evolutionary dynamics of urinary pheromone signal production in *Mus* reveals three key findings. First, our results indicate that the information content of MUPs has changed over time. More basal species have single MUPs expressed by males, though the identity of those proteins varies. In more recently derived lineages, both males and females express MUPs though the relative differences in expression patterns vary across species. Second, variation in relative expression between males and females indicates that the information content of a given protein changes across lineages. Specifically, the male specificity of *Mup3* and *Mup20* varies across species, suggesting that the responses to these proteins are also likely to differ across taxa. Third, the extreme diversity in central MUP genes in house mice is specific to the *M. musculus* species group and predates subspecies diversification. Moreover, most proteins in our dataset are relatively rare across samples and only a few are shared across subspecies, suggesting that protein diversity evolves and turns over rapidly across house mouse populations. Notably, the result that diversity is overrepresented in sites with greater exterior exposure that may interact with vomeronasal receptors is consistent with the functional diversification of MUPs as pheromone signaling molecules.

### Two modes of pheromone evolution in mouse urinary scent marks

Pheromone blends can evolve through changes in the relative ratios of semiochemicals or via changes to the specific pheromone compounds (Symonds and Elgar 2008; Wyatt 2010). Both modes of pheromone evolution appear to occur in mice. Variation in the identity of MUPs expressed in the liver has been achieved through two different mechanisms: gene family expansion and changes in patterns of gene expression.

There appear to have been multiple gene family expansions followed by sequence divergence contributing to differences in MUP blends across species. At least four gene expansions are well-supported by present data. However, additional expansions seem likely; better assembled sequences of the entire *Mup* gene cluster across species will provide a clearer picture of gene family evolution. Previous examinations of *Mup* gene family evolution have indicated an independent expansion within *Mus* relative to *Rattus* (Logan et al. 2008). The present analysis provides further details on the relative timing and nature of at least four expansion events (Figs 2, 4, S1).

1. At least one initial expansion event appears to have occurred relatively early within *Mus*, as indicated by the diversity of genes present in the *M. pahari* genome and the fact that *M. mattheyi* and *M. pahari* appear to utilize different *Mup* genes as pheromones.
2. An expansion of *Mup3*-like genes appears to have occurred on the lineage leading to *M. caroli* that is absent in other species.
3. An initial expansion of central *Mup* genes occurred in the common ancestor of *M. spretus, M. spicilegus, M. macedonicus* and *M. musculus.* All species appear to have at least 3 central *Mup* genes though the genes do not fall into 3 clades with orthologs for each species. Previous studies examining patterns of diversity among central *Mup* genes in a wild population of *M. m. domesticus* found evidence of gene conversion (Sheehan et al. 2016). Given the similarity in the gene sequences among central *Mup* genes within each species, gene conversion may obscure some aspects of the timing of diversification within and between species.
4. Prior to the diversification of the three subspecies of house mouse approximately 350,000 years ago there seems to have been an additional expansion of central *Mup* genes. This expansion has produced individually distinctive MUP pheromone blends used for individual recognition (Hurst et al. 2001; Kaur et al. 2014).

Additional variation in urinary pheromone blends has been achieved through differential expression of genes across species. Previous work has demonstrated that within a population of *M. m. domesticus* there is variation in whether or not mice express certain central *Mup* genes (Sheehan et al. 2016). The present study demonstrates that different species of mice typically possess copies of central MUPs, *Mup3* and *Mup20* in their genome that are not expressed in the liver (Fig 3, Table 1). Why are non-expressed genes maintained in species’ genomes? We only examined genes predicted to produce full-length proteins suggesting they are not pseudogenes. Of course, a lack of expression in the liver does not mean the *Mup* genes are not expressed in other tissues. There is evidence that MUP proteins are excreted in bodily fluids other than urine and may serve a signaling function in other secretions (Stopková et al. 2016; Černá et al. 2017). Thus, selective pressures on the maintenance and diversification of *Mup* genes is likely driven by social communication not limited of territory marking. Moreover, *Mup3, Mup20* and *Mup11* are all expressed in tissues other than the liver (Finger et al. 2016) and may have other yet to be described functions that may favor the maintenance of these genes in the genome.

### Variation in sex-specificity of MUP3, a presumed ‘male-specific’ pheromone

Previous studies have shown that expression of MUP3 is male-limited in the lab strain C57BL/6J (aka B6), leading to the assumption that this is a male-specific protein (Mudge et al. 2008). Behavioral work in B6 has further shown that MUP3 is sufficient to induce males to countermark and promotes aggressive behavior in the presence of another male (Kaur et al. 2014), as might be expected for a pheromone that signals male ownership of a scent mark. However, we find robust evidence for female expression of *Mup3* in *M. m. domesticus*, *M. m. castaneus, M. macedonicus*, and of Mup3-like genes in *M. caroli*. While overall *Mup* gene expression is low in female *M. macedonicus, Mup3* genes are expressed at relatively high levels compared to central *Mup* genes. Whether expression leads to biologically relevant excretion of Mup3 proteins in female *M. macedonicus* remains to be determined. Female *M. m. domesticus* and *M. m. castaneus*, however, do express reasonably high quantities of *Mup3*, such that the protein would likely be a relevant part of their urinary scent. This revelation is especially surprising since many wild-caught *M. m. domesticus* female urine samples have been analyzed but MUP3 has not been detected (e.g. Hurst et al. 2017). However, MUP3 does not show up on standard analyses of urine protein content using mass spectrometry due to glycosylation that increases the mass, though it is detectable via other methods (discussed further as B6 gene18 in Mudge et al. 2008). Gel electrophoresis detects expression in male B6 urine but not female. Thus, the peculiar features of Mup3 and a dearth of transcriptomic analyses have likely led to a misestimation of its typical expression patterns in wild populations.

Comparative analyses of pheromones in *Drosophila* show a similar pattern of variation in sex specificity of key compounds among species (Seeholzer et al. 2018). Changes in sex-limited expression of key pheromones may be a straightforward way for species to diverge in sexual signaling from close relatives where hybridization may be disfavored, as is the case among mouse subspecies (Smadja and Ganem 2002; Smadja et al. 2004). While these results do not call into question previous behavioral studies on the effects of MUP3 on male behavior, they do suggest the possibility that neural mechanisms for processing the same pheromone compounds have diverged among closely related species of mice, as has been show in *Drosophila* (Seeholzer et al. 2018). Repeating studies of *Mup3* effects on behavior in wild-derived strains or wild-caught mice from different subspecies will be important to understanding the behavioral selection pressures influencing the evolution of urinary signals in mice.

### Liver expression of darcin is a recent phenomenon

Darcin (Mup20) is a male-biased protein that is highly expressed in territorial male *M. m. domesticus* (Roberts et al. 2010) and *M. m. musculus* (Thoß et al. 2019). Our data support these previous studies. We detect very minimal expression in male *M. spretus,* and do not detect darcin expression in *M. m. castaneus, M. macedonicus* or *M. spicilegus*. Behavioral and neurobiological studies have demonstrated that darcin promotes female *M. m. domesticus* attraction to male scents and memory for the location where those scents were encountered (Roberts et al. 2012). Furthermore, darcin leads to increased countermarking in B6 males (Kaur et al. 2014). Based on patterns of gene expression, we predict that *M. m. musculus* would respond similarly to darcin, whereas response would be different or absent in other closely related species. Recent changes in expression of darcin among closely related species provides an excellent opportunity to examine the mechanisms of species differences in responses to pheromone signals. This is especially true given the rich set of responses documented to darcin (Roberts et al. 2010; Roberts et al. 2012; Kaur et al. 2014; Hoffman et al. 2015).

### Individual identity signatures and patterns of pheromone diversification

Scent signatures are commonly used for individual recognition across diverse animal taxa including ants (D’Ettorre and Heinze 2005), fish (Thünken et al. 2009), lizards (Carazo et al. 2008), and many mammals (Johnston 2003; Thom and Hurst 2004). Traits used for individual recognition are expected to be under negative frequency-dependent selection (Dale et al. 2001; Tibbetts et al. 2017), as this process will maintain diversity in traits needed for differentiating among individuals. Despite wide interest in recognition abilities, little work has examined the molecular diversity and evolution of traits used for recognition, likely because the underlying molecular basis is typically unknown. Because they are directly derived from gene products, MUPs provide an unusual opportunity to investigate the evolution of individual identity. The present study provides key insights in the origins of individual identity and the processes shaping trait diversification.

Our data refine the estimate for the expansion of central MUPs that mediate individual recognition in house mice. Previous work had suggested that central MUP diversification may date to the split between *M. musculus* and *M. spretus* (Mudge et al. 2008) while other authors had suggested high population density associated with human commensalism may have driven the expansion of central *Mup* genes (Logan et al. 2008). The present study indicates that there were two distinct phases of central MUP expansion: a minor expansion in the common ancestor of *M. musculus* and *M. spretus* and a second larger expansion prior to the diversification of *M. musculus* subspecies, which occurred approx. 350,000 years ago (Geraldes et al. 2008). Thus, the dramatic expansion of central MUPs that mediate recognition is more recent than previously estimated but predates commensalism with humans, because commensalism likely evolved independently in the three subspecies. The ability to advertise individual identity in urinary scent marks is likely to be beneficial in dense populations observed in *M. musculus* both in feral and commensal settings (Hurst 1987; Pocock et al. 2004). One tantalizing possibility is that increased identity information in urinary signals facilitated the evolution of commensalism with humans approximately 10,000 years ago. Identity signals may have facilitated the independent evolution of commensalism in each subspecies. High population densities associated with human commensalism in mice should increase interaction rates among animals, making identity information a potentially vital mediator of territory marks. A correlation between social structure and the extent of identity has also been reported for alarm calls in marmots (Pollard and Blumstein 2011), indicating that social interactions can favor increased signal individuality. While it is likely that commensalism has further selected for the elaboration and diversification of central *Mup* genes, the data presented here (Fig 4) indicate that the initial elaboration happened prior to subspecies diversification.

Analyses of the MUP isoforms found across populations and subspecies of house mice provide insights into the dynamics of selection on individual identity signaling phenotypes. Consistent with a model of negative frequency-dependent selection favoring novel proteins, we find evidence for rapid turnover of protein composition across *M. musculus* subspecies and populations. Of 77 distinct proteins, only 2 proteins were shared across all subspecies, and 50 proteins were specific to a single sample (though not necessarily a single individual as some samples in the NCBI Short Read Archive were pools of wild-caught individuals). Given the relatively small sample of mice studied, further sampling is almost certain to uncover many more central MUP isoforms in wild mouse populations.

Compared to rarity of shared proteins across populations, amino acid substitutions are more commonly shared among proteins. Amino acid variants on the exterior of the protein are much more likely to be found in a greater number of protein variants. The existence of a relatively small number of hyper-variable sites has two implications for the evolution of MUP protein diversity. First, the observation is consistent with previous analyses suggesting that gene conversion is likely commonplace among central MUPs (Mudge et al. 2008; Sheehan et al. 2016), as they do not segregate into discrete groups but display a mosaic distribution across proteins (Fig 5a). Second, these few highly variable sites are likely candidates for interaction sites between central MUP ligands and their corresponding vomeronasal receptors. Thus, the sharing of a modest number of amino acid variants across proteins may be the result of convergent evolution shaped by a limited set of possible amino acid variants.

### The utility of wild house mice for understanding pheromone function and evolution

The rapid expansion of the neurobiological toolkit in lab mice for assessing how the neural processing of pheromones can be readily translated to wild house mice and their relatives. The diversity of predicted pheromone blends documented in this study suggests that wild mice and their relatives will be an unusually valuable system for understanding the evolution of complex pheromone-mediated behaviors. Further research is now needed to understand how mice differ in the use of urinary scent marks as well as how they respond to individual components of scent blends. Coupled with the toolset available for mouse genetics and neurobiology, studies of wild mouse scent marks have the potential to substantially further our understanding the role of pheromone signals in behavior and evolution.

## ACKNOWLEDGEMENTS

Funding for this research was provided by Cornell University. The *M. macedonicus* XBS and *M. spicilegus* ZRU strains were originally developed by the Wild Mouse Genetic Repository (University of Montpellier). We thank Tory Hendry for assistance with generating gene trees.

## LIST OF SUPPLEMENTAL FILES

1. Supplemental File1: contains S2, S3 and details related to Fig 5.

2. Supplemental File2: Coding sequences of *Mup* genes expressed in the liver

3. Supplemental File3: Coding sequences of *Mup* genes from BLAST search of genomes

4. Supplemental File4: Tree file of liver expressed gene sequences

**Figure S1.**
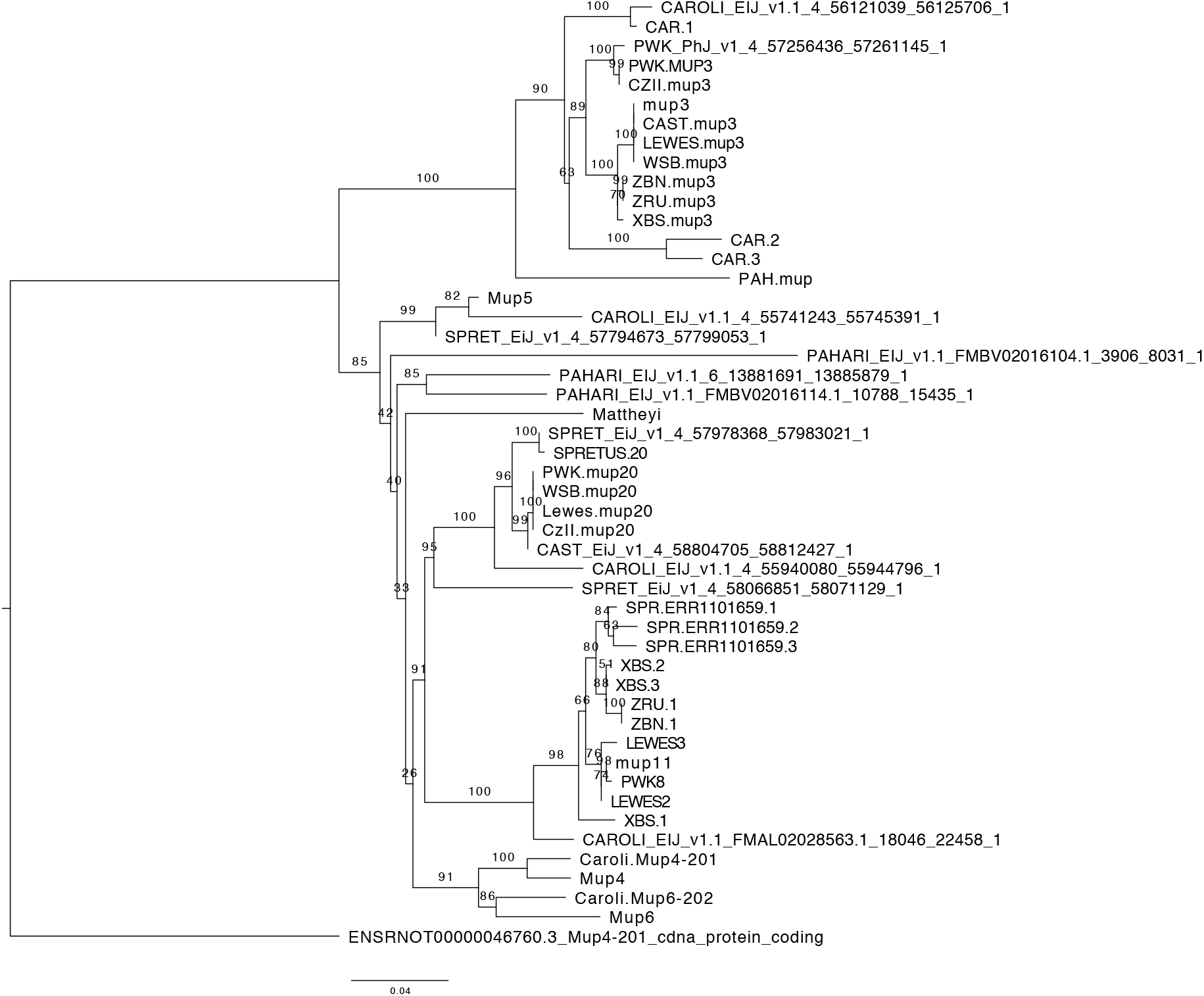
Gene tree of expressed and non-expressed *Mup* gene sequences. The ML gene tree shows *Mup* gene sequenced identified from transcriptomes as well as predicting coding sequences identified from BLAST searches again published genomes. Sequences from genomes are given as SpeciesGenomeName_Version_Scaffold_StartBase_EndBase. The scale bar shows substitutions per site. All bootstrap values are shown.

